# Changes in Motor Unit Activity of Co-activated Muscles During Dynamic Force Field Adaptation

**DOI:** 10.1101/2025.03.11.642567

**Authors:** Yori R. Escalante, Shancheng Bao, Yuming Lei

## Abstract

Muscle co-contraction plays a critical role in motor adaptation by minimizing movement errors and enhancing joint stability in novel dynamic environments. However, the underlying changes in motor unit (MU) activity within co-activated muscles during adaptation remain largely unexplored. To investigate this, we employed advanced electromyography sensor arrays and signal processing to examine MU activation in the triceps brachii (agonist) and biceps brachii (antagonist) during a reaching task under force-field perturbation. Our results revealed a gradual reduction in movement errors and an increase in velocity with adaptation, accompanied by a decrease in muscle co-contraction from early to late adaptation phases. This reduction was primarily driven by increased triceps activity, while biceps activity remained unchanged throughout the adaptation process. At the MU level, recruitment, amplitude, and firing rate increased in both muscles during adaptation compared to baseline (without force-field perturbation). However, from early to late adaptation phases, triceps MU amplitude continued to increase, while its firing rate stabilized, suggesting a shift in force generation strategy. In contrast, biceps MU activity remained stable throughout the adaptation. These findings indicate that the reduction in co-contraction during motor adaptation is likely mediated by a shift in motor unit control strategy within the agonist muscle. The increased reliance on MU amplitude modulation rather than firing rate in later adaptation may represent a mechanism for optimizing force production while maintaining movement accuracy and joint stability in dynamic environments.

**NEW & NOTEWORTHY:** This study examines how motor unit (MU) activity changes during motor adaptation in dynamic environments. We show that reduced co-contraction during adaptation is primarily driven by increased agonist MU amplitude rather than firing rate changes. In contrast, antagonist MU activity remains stable. These findings highlight a shift in MU control strategy that optimizes force production while maintaining movement accuracy, providing new insights into the underlying neuromuscular mechanisms of motor adaptation.

## Introduction

In an ever-changing and unpredictable world, the ability to adapt motor actions is essential for maintaining stability and effectively interacting with the environment. Whether navigating uneven terrain (1) or manipulating objects with varying weight and texture (2), the central nervous system (CNS) continuously refines movement strategies to optimize efficiency and accuracy in response to external forces (3). This adaptability is governed by two key control strategies. One strategy involves learning internal models that predict the sensory consequences of motor commands and using prediction errors to generate appropriate forces (4, 5). Another strategy relies on impedance control, where muscle co-contraction adjusts limb and joint stiffness to improve movement stability and accuracy (6, 7).

Muscle co-contraction is critical for dynamic adaptation by stabilizing joints and compensating for inaccuracies in the internal model when exposed to unpredictable perturbations (6, 8). Furthermore, co-contraction, by stiffening the limb, effectively minimizes kinematic errors and ensures more controlled, predictable movements during adaptation to novel force perturbations (7, 9). As adaptation progresses and internal model predictions become accurate, muscle co-contraction is gradually reduced (10, 11). This reduction marks a shift toward a more efficient motor control strategy, characterized by enhanced movement precision and likely lower energy expenditure (7, 11, 12). Nevertheless, a certain level of co-contraction remains crucial for maintaining joint stability, particularly in environments with high variability or external disturbances (7, 11). By dynamically regulating co-contraction, the CNS optimally balances movement efficiency and stability, ensuring adaptability while maintaining robustness against unexpected perturbations (9).

Muscle co-contraction during dynamic adaptation has been examined through psychophysical and neuroimaging methodologies (6, 12–15). For instance, psychophysical studies suggest that elevated co-contraction characterizes the early adaptation phase, increasing limb stiffness to counteract unpredictable perturbations. As adaptation progresses, co-contraction gradually decreases. In the late adaptation phase, the CNS fine-tunes co-contraction levels to enhance movement efficiency while minimizing metabolic costs (6). Neuroimaging studies implicate the cerebellum as a key neural substrate for the dynamic regulation of muscle co-contraction. During early adaptation, increased functional connectivity between the cerebellum and the inferior parietal lobule (IPL) suggests that this cortico-cerebellar loop is critical for initial co-contraction modulation (15). In contrast, the reduction in co-contraction observed in the late adaptation phase is associated with a distinct neural network involving the cerebellum, superior frontal gyrus (SFG), and motor cortical regions, indicating a transition in neural control mechanisms as adaptation becomes more refined (15).

Despite these insights, the underlying changes in motor unit (MU) activity within co-activated muscles during dynamic adaptation remain largely unexplored. MUs are the fundamental units of muscle force generation, and their recruitment and modulation are critical for precise movement control (16–18). We hypothesize that MU activity will differ across adaptation phases, reflecting changes in neuromuscular control strategies. During early adaptation, increased muscle co-contraction will be associated with distinct MU recruitment patterns in both agonist and antagonist muscles to enhance joint stability. As adaptation progresses, we expect a shift in MU control, leading to adjustments in activation patterns that optimize movement efficiency and stability. To test this hypothesis, we will extend our previous work using advanced electromyography (EMG) sensor arrays and signal processing techniques to decode MU activity (19–21). This study aims to characterize how MU recruitment, firing patterns, and amplitude modulation evolve throughout adaptation, providing new mechanistic insights into how the CNS optimizes movement stability, accuracy, and efficiency in response to dynamic environments.

## Methods

### Participants

The study recruited 17 right-handed, healthy participants aged 18–30, all of whom were naive to the experimental paradigm and study objectives. The research protocols were reviewed and approved by the Institutional Review Board of Texas A&M University. Prior to participation, all individuals provided written informed consent, in compliance with ethical guidelines set by the local ethics committee at Texas A&M University and in accordance with the Declaration of Helsinki.

### EMG recording

Electromyographic (EMG) activity was recorded from the biceps brachii and triceps brachii muscles utilizing two Trigno Galileo surface EMG sensor arrays (Delsys, Inc., MA, USA) positioned over the muscle belly of each. Prior to sensor application, the participants’ skin at the designated electrode sites was prepared by cleaning with an abrasive gel followed by an alcohol solution to ensure optimal electrode-skin contact and minimize impedance for high-quality EMG recordings. Each sensor array consisted of four cylindrical probes, allowing for multi-channel signal acquisition. EMG signals were derived through pairwise differentiation of the four electrodes within each array, resulting in four distinct channels per sensor. These signals were then amplified, digitized at a sampling rate of 2222 Hz, and stored digitally on a computer for subsequent offline analysis.

### Dynamic Adaptation Task

The dynamic adaptation task in this study involved an abrupt force-field adaptation paradigm using a bilateral robotic exoskeleton, the KINARM (BKIN Technologies Ltd, Kingston, ON, Canada). Participants were seated in the KINARM’s built-in chair, facing a testing table, with both arms positioned and supported by the device’s mechanical arms. The KINARM was equipped with a virtual reality system that projected a feedback cursor (representing the index fingertip) and a target circle onto a horizontal display, ensuring that all visual feedback appeared in the same plane as the participant’s arm. To eliminate direct visual feedback of their hands and arms, a black occlusion cover was placed beneath the display. Participants performed a series of goal-directed reaching movements using their right arm, controlling the feedback cursor from a start circle to a target circle displayed on the screen (Fig. 1A). Both circles measured 1 cm in radius and were positioned 10 cm apart. At the start of each trial, participants aligned the cursor with the center of the start circle. After a 500-ms delay, the target circle appeared, cueing the participant to initiate movement. They were instructed to reach the target quickly and accurately. No trials were excluded based on movement speed, allowing for natural variability in response times. Throughout the task, participants received continuous visual feedback of their hand position, represented by a white cursor. Upon reaching the target, the robotic exoskeleton passively returned the participant’s arm to the start position to maintain consistency across trials. During the dynamic adaptation phase, participants performed reaching movements while experiencing externally applied force perturbations generated by the KINARM exoskeleton. Specifically, a velocity-dependent force of 15 N was applied leftward, perpendicular to the reaching direction, while a 30 N viscous force was imposed forward, opposing the movement (Fig. 1A). The leftward force depended on movement velocity, meaning that as participants moved faster, the perturbation magnitude increased, requiring them to adjust their trajectory to counteract the lateral deviation. The forward viscous force introduced resistance proportional to velocity, mimicking the sensation of moving through a thick fluid, demanding greater effort to maintain speed and accuracy.

**Figure 1.**
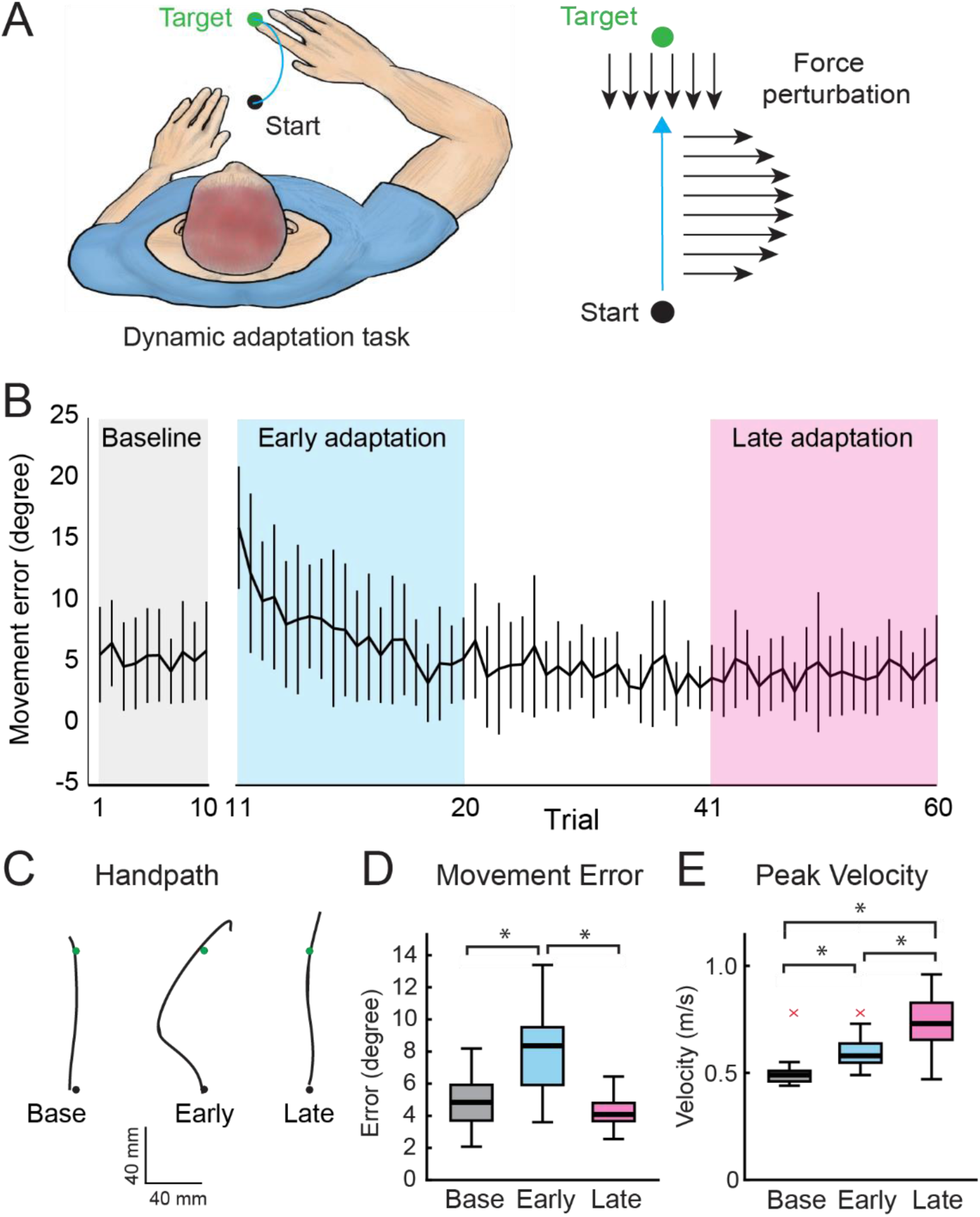
(**A**) Schematic of the force-field adaptation task, where subjects performed reaching movements in both a null and perturbed space. Perturbations included a 15 N velocity-dependent force applied leftward and a 30 N viscous force opposing movement. (**B**) Directional errors across baseline (grey), early adaptation (blue), and late adaptation (red) trials. (**C**) Representative hand paths for baseline, early adaptation, and late adaptation trials. (**D**) Box plots illustrating significant differences in movement error between baseline and early adaptation trials, as well as between early and late adaptation trials, with no significant differences observed between baseline and late adaptation trials. (**E**) Box plots showing significant differences in movement velocity across all trial comparisons: baseline vs. early adaptation, early vs. late adaptation, and baseline vs. late adaptation. “x” denotes outliers—data points that fall significantly outside the expected range.

### Experimental Paradigm

Participants were seated in the KINARM’s built-in chair, with seat height and arm supports adjusted for comfort and optimal task performance. A calibration process aligned the participant’s arm with the robotic arm to ensure accurate reaching movements. Once fitting and calibration were complete, wireless EMG electrodes were placed. For the biceps brachii, the participant performed elbow flexion against resistance to locate the muscle belly, where the electrode was positioned parallel to the muscle fibers. For the triceps brachii, elbow extension from 90° to 180° against resistance identified the muscle belly for electrode placement. Electrodes were securely attached and positioned to minimize movement artifacts. The experiment consisted of two blocks: a 20-trial baseline block and a 60-trial adaptation block. During baseline, no external forces were applied, and participants reached the target as quickly and accurately as possible. In the adaptation block, force perturbations were introduced, including a 15 N velocity-dependent force applied leftward and a 30 N viscous force opposing movement. Despite these perturbations, participants were instructed to maintain speed and accuracy. The adaptation block was further divided into three phases to assess learning: Early adaptation (trials 1–20), Middle adaptation (trials 21–40), and Late adaptation (trials 41–60). Kinematic and EMG data were synchronized to ensure precise alignment between movement execution and muscle activity recordings.

### Kinematics and EMG Activity

The hand’s position in the X-Y coordinate system was recorded at 1000 Hz and processed using a 15 Hz low-pass filter to minimize high-frequency noise. Movement velocity was computed as the displacement between consecutive positions per unit time. Peak velocity was identified as the highest instantaneous velocity within each movement trial by analyzing the velocity-time profile. To assess performance accuracy, direction error (DE) was calculated as the angular difference between two vectors: one extending from the start circle to the target and the other from the hand position at movement onset to its position at peak velocity. Kinematic and EMG data were synchronized to ensure precise temporal alignment between movement execution and muscle activity recordings. For EMG processing, the mean rectified EMG activity was smoothed using a Gaussian smoothing kernel to reduce signal fluctuations. RMS EMG was calculated using a 100 ms moving-average window across the entire reaching movement in each trial to quantify the magnitude of muscle activation. Muscle co-contraction levels were assessed by computing RMS EMG for biceps and triceps muscle pairs, capturing the degree of simultaneous activation.

### EMG Signal Decomposition

The recorded EMG signals were decomposed into distinguishable motor units (MUs) using the decomposition algorithm developed by De Luca (22–24), with the process conducted separately for baseline (20 trials), early adaptation (20 trials), middle adaptation (20 trials), and late adaptation (20 trials) to assess changes in motor unit behavior across sessions. For each identified MU, the algorithm provided motor unit discharge times and motor unit action potential (MUAP) templates. To extract MU waveforms, including amplitude and shape, spike-triggered averaging (STA) was applied to the raw surface EMG signals, using the MU discharge times as trigger points for STA calculation. To ensure the reliability of the STA estimate (19, 25), two validation tests were performed: the coefficient of variation (CV) of the MUAP amplitude and the maximum linear correlation coefficient between the STA estimate and the decomposition-estimated MUAP templates. MUs with a correlation coefficient greater than 0.7 and a CV of MUAP amplitude below 0.3 were retained. For each retained MU, the MUAP amplitude of the STA template was used as an estimate of MU amplitude, measured as the voltage difference between the minimum and maximum peaks. Additionally, the MU discharge rate was calculated over the entire session duration.

### Statistical analysis

To assess the effect of SESSION (Baseline, Early Adaptation, Late Adaptation) on kinematics, EMG activity, and motor unit activity, Repeated-Measures ANOVAs were conducted. These analyses examined the effect of SESSION on movement error and peak velocity, providing insights into motor performance changes across adaptation phases. Additionally, a Repeated-Measures ANOVA with SESSION and MUSCLE (Biceps and Triceps) as within-subject factors was performed to evaluate changes in EMG activity magnitude over time and between muscles. To analyze muscle co-contraction patterns, a Repeated-Measures ANOVA was conducted to compare co-contraction levels across sessions. Additionally, Repeated-Measures ANOVAs were used to determine whether SESSION and MUSCLE had a significant effect on motor unit (MU) properties, including MU number, MUAP amplitude, and MU discharge rate, which reflect changes in neural drive and MU recruitment throughout the adaptation process. The Shapiro-Wilk test was used to evaluate whether the data followed a normal distribution, while Levene’s test for equality of variances assessed the homogeneity of variances across conditions. Mauchly’s test for sphericity was conducted to determine whether the assumption of equal variance in repeated-measures data was met. If the data violated the assumption of normality, a log transformation was applied to approximate a normal distribution. In cases where the assumption of sphericity was not satisfied, the Greenhouse-Geisser correction was applied to adjust the degrees of freedom, ensuring accurate statistical inferences. All group data are reported as mean ± standard deviation (SD) throughout the text.

## Results

### Kinematics

During the baseline and adaptation sessions, all participants performed point-to-point reaching movements with their right arm, with no force perturbations applied during baseline and force perturbations introduced during adaptation. Figure 1B depicts the variations in direction error (DE) across trials during both the baseline and adaptation sessions. As anticipated, DEs remained minimal during the baseline phase, indicating precise reaching in the absence of perturbations. Upon introduction to the novel force perturbation, DEs increased significantly, reflecting initial movement disruptions. However, DEs progressively diminished throughout the adaptation session, demonstrating gradual motor adaptation. Figure 1C illustrates the typical hand path of a representative participant during baseline, early adaptation, and late adaptation. During baseline, the hand path remains relatively straight, indicating unperturbed reaching movements. In early adaptation, the hand path initially deviates leftward due to the application of a velocity-dependent force in that direction, reflecting the participant’s initial response to the perturbation. By late adaptation, the hand path is once again relatively straight, suggesting a significant degree of motor adaptation and successful compensation for the applied force. A Repeated-Measures ANOVA revealed a significant effect of SESSION on DE (F_(2, 32)_ = 21.2, p < 0.001). DEs during baseline were 5.02 ± 1.77°. Upon introduction to the force perturbations, DEs increased significantly to 8.05 ± 2.69° during early adaptation (p < 0.001; Fig. 1D). However, DEs progressively decreased to 4.17 ± 0.95° by late adaptation, demonstrating the participants’ ability to adjust to the leftward force perturbation (p < 0.001). A Repeated-Measures ANOVA also revealed a significant effect of SESSION on peak velocity (F_(2, 32)_ = 30.3, p < 0.001). Peak velocity increased during late adaptation (0.73 ± 0.13 m/s, p < 0.001) compared to early adaptation (0.60 ± 0.08 m/s; Fig. 1E), indicating an adaptation to the viscous force opposing movement.

### EMG Activity

We analyzed the activation magnitude of the biceps and triceps muscles during baseline and adaptation sessions. A Repeated-Measures ANOVA revealed a significant effect of SESSION (F_(2,32)_ = 32.1, p < 0.001), FORCE (F_(1,16)_ = 7.5, p = 0.015), and their interaction (F_(2,32)_ = 12.0, p < 0.001) on EMG activity magnitude. Post hoc analysis showed that biceps EMG activity increased significantly during early adaptation compared to baseline (p < 0.001) but remained unchanged between early and late adaptation (p = 0.2; Fig. 2A). In contrast, triceps EMG activity also increased significantly from baseline to early adaptation (p < 0.001) and continued to increase from early to late adaptation (p < 0.001; Fig. 2B), indicating progressive adjustments in triceps activation as adaptation progressed. A Repeated-Measures ANOVA revealed a significant effect of SESSION (F_(2,32)_ = 15.7, p < 0.001) on biceps and triceps co-contraction levels. Post hoc analysis showed that the co-contraction level during early adaptation was 0.87 ± 0.35, but it significantly decreased in late adaptation (0.65 ± 0.18, p = 0.006; Fig. 2C), suggesting a reduction in co-contraction as participants adapted to the force perturbation. A co-contraction value close to 1 indicates balanced activation between the biceps and triceps, whereas values greater than 1 suggest greater biceps activation and values less than 1 indicate greater triceps activation.

**Figure 2.**
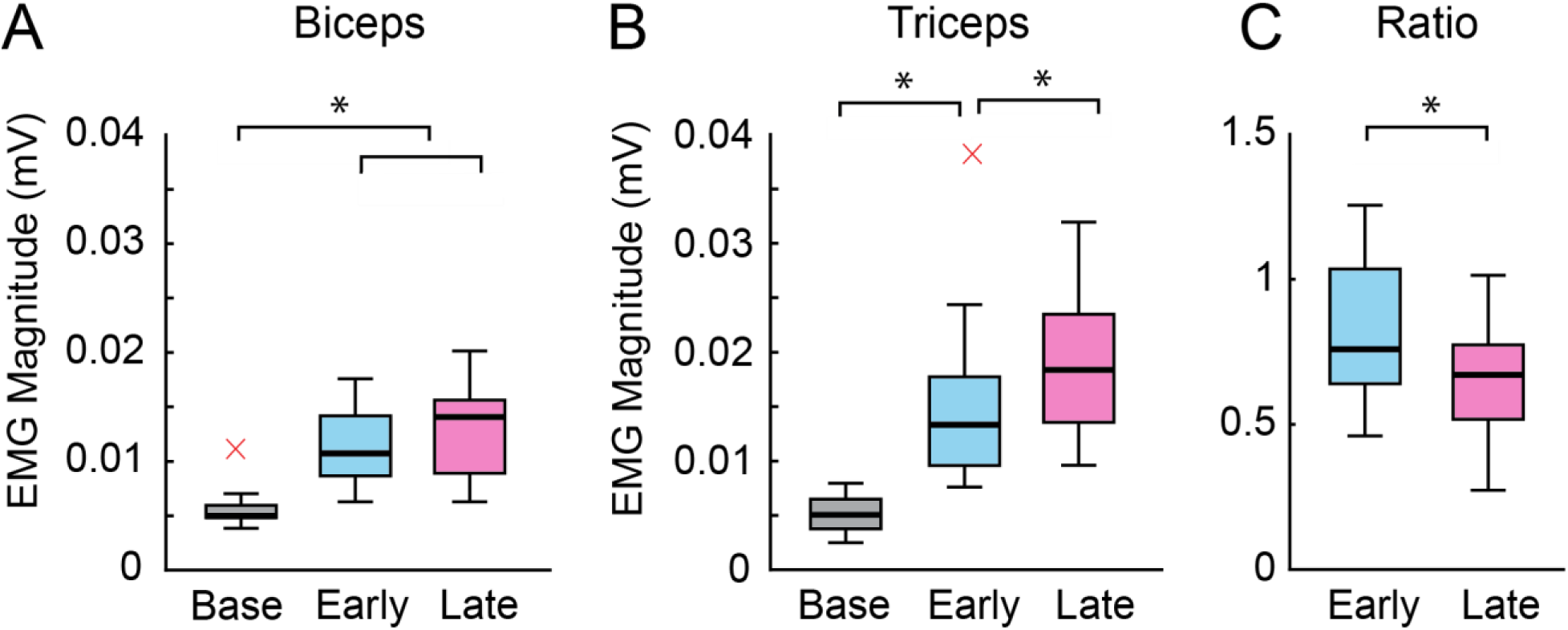
EMG magnitude and muscle co-contraction across trials. (**A**) EMG magnitude of the biceps brachii during baseline, early adaptation, and late adaptation trials, with significant differences observed between baseline and both adaptation phases. (**B**) EMG magnitude of the triceps brachii across the same trial conditions, showing significant differences between baseline and both early and late adaptation trials, as well as between early and late adaptation trials. (**C**) Co-contraction ratio between the biceps brachii and triceps brachii, with a significant difference detected between early and late adaptation trials. “x” denotes outliers—data points that fall significantly outside the expected range.

### Motor Unit Activity

A total of 373 motor units (MUs) were decomposed from the biceps muscle and 384 MUs from the triceps muscle. During the baseline session, reliable MUs could not be decomposed for four participants in the biceps and one participant in the triceps, likely due to low muscle activity during reaching movements in the absence of force perturbations. However, during the adaptation session, reliable MUs were successfully detected in all participants. A Repeated-Measures ANOVA revealed a significant effect of SESSION (F_(2,24)_ = 7.2, p = 0.003) on MU number, while FORCE (F_(1,12)_ = 0.03, p = 0.9) and the SESSION × FORCE interaction (F_(2,24)_ = 1.6, p = 0.2) showed no significant effects. Significantly more MUs were decomposed during the adaptation session than in the baseline session for both the biceps and triceps (p < 0.02; Fig. 3A-B). However, there was no significant difference in MU count between early and late adaptation (p > 0.8; Fig. 3A-B) in either muscle.

**Figure 3.**
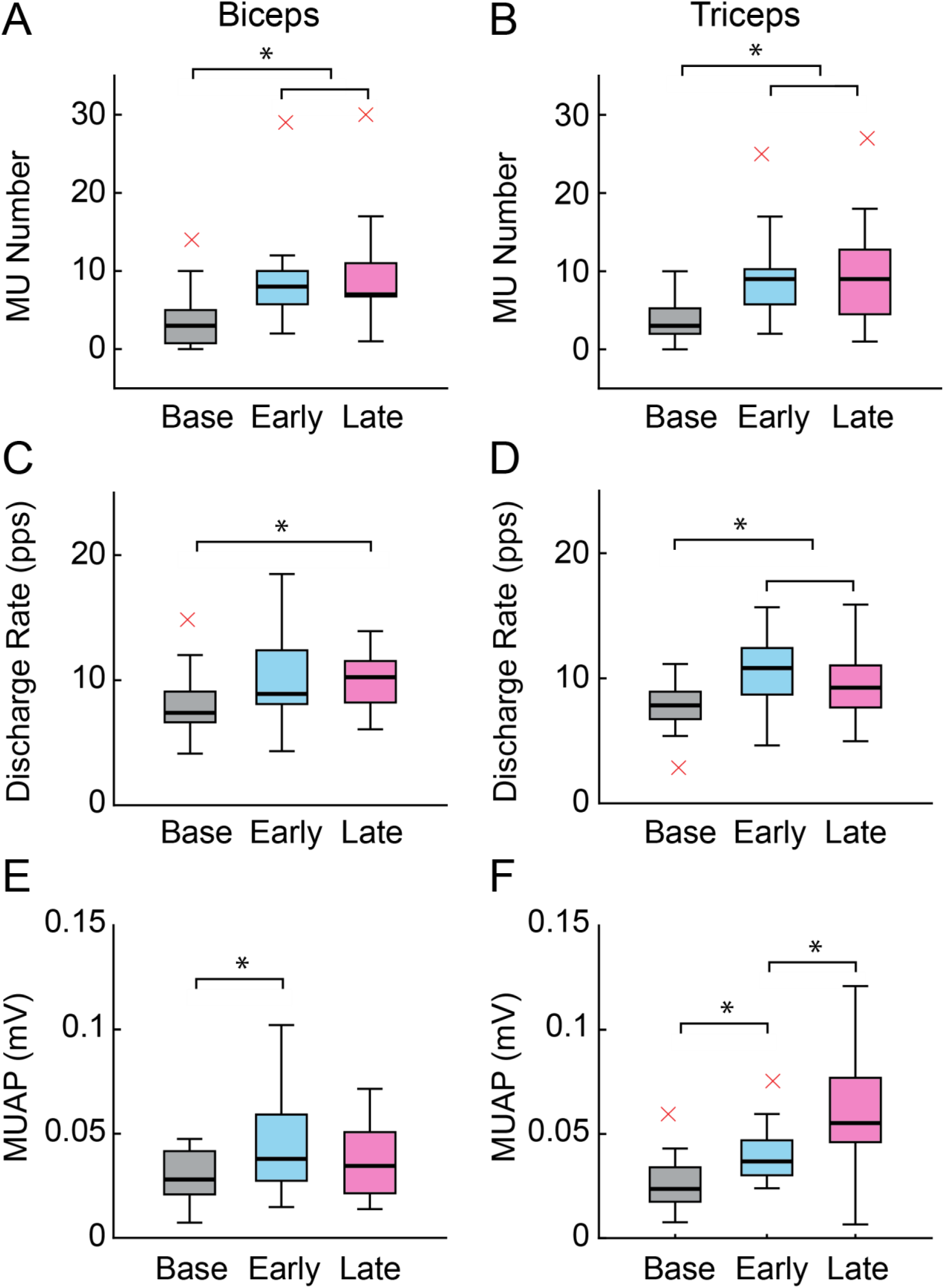
Changes in motor unit (MU) characteristics across baseline, early adaptation, and late adaptation trials. (**A, B**) MU number significantly differed between baseline and both early and late adaptation trials in the biceps brachii (A) and triceps brachii (B). (**C, D**) MU discharge rate showed significant differences between baseline and late adaptation in the biceps brachii (C) and between baseline and both early and late adaptation trials in the triceps brachii (D). (**E, F**) MU action potentials (MUAP) differed significantly between baseline and early adaptation in both the biceps brachii (E) and triceps brachii (F), as well as between early and late adaptation trials in the triceps brachii (F).

Figure 3C-D illustrates MU firing rates across baseline and adaptation sessions. A Repeated-Measures ANOVA revealed a significant effect of SESSION (F(2,24) = 7.8, p = 0.002) on MU firing rate, while FORCE (F(1,12) = 0.9, p = 0.3) and the SESSION × FORCE interaction (F(2,24) = 0.6, p = 0.5) showed no significant effects. In the biceps muscle, MU firing rate significantly increased during late adaptation compared to baseline (p = 0.04), with no significant difference between early and late adaptation (p = 0.8). In the triceps muscle, MU firing rate was significantly higher during both early and late adaptation compared to baseline (p < 0.04), though there was no significant difference between early and late adaptation (p = 0.3).

Figure 3E-F illustrates MUAP amplitude across baseline and adaptation sessions. A Repeated-Measures ANOVA revealed a significant effect of SESSION (F(2,24) = 14.5, p < 0.001) and a SESSION × FORCE interaction (F(2,24) = 7.2, p = 0.003) on MUAP amplitude, while FORCE (F(1,12) = 0.9, p = 0.3) showed no significant effects. In the biceps muscle, MUAP amplitude was significantly higher during both early and late adaptation compared to baseline (p < 0.02), with no significant difference between early and late adaptation (p = 0.2). However, in the triceps muscle, MUAP amplitude was significantly higher during both early and late adaptation compared to baseline (p < 0.005) and continued to increase from early to late adaptation (p = 0.002), emphasizing a progressive change in MU recruitment throughout adaptation.

## Discussion

This study investigated motor unit (MU) activity in the co-activated triceps and biceps brachii during reaching movements in a novel force field. Our findings reveal distinct changes in MU recruitment, firing patterns, and amplitude throughout adaptation. Initial force-field exposure led to increased EMG activation magnitude in both muscles and heightened co-contraction, likely serving to minimize initial movement errors and stabilize the limb in the novel dynamic environment. As adaptation progressed, movement errors decreased, velocity increased, and muscle activity patterns evolved. Co-contraction diminished from early to late adaptation, driven primarily by increased agonist triceps activity, while antagonist biceps activity remained relatively stable. Crucially, our MU-level analysis revealed that while both muscles initially exhibited increased recruitment, amplitude, and firing rates compared to baseline, the transition to late adaptation was characterized by a selective increase in triceps MU amplitude and a stabilization of its firing rate. In contrast, biceps MU activity remained unchanged throughout adaptation. These findings demonstrate a neural strategy that initially employs broad muscle activation to reduce errors, followed by a transition to a more efficient force production mechanism. The increased reliance on triceps MU amplitude modulation, rather than firing rate, suggests an optimization of force output for fast and accurate movement in dynamic environments. This shift in agonist motor unit control appears to be the key mechanism underlying reduced co-contraction, highlighting an efficient adaptive process within the motor system.

Previous studies demonstrated that when faced with unfamiliar forces or dynamics, the CNS initially increases agonist-antagonist co-contraction to stabilize the limb, then gradually reduces co-contraction as adaptation progresses. For example, Franklin et al. (2003) showed that early adaptation is characterized by elevated co-contraction, which stiffens the arm against unpredictable disturbances (6). As performance errors decrease, co-contraction progressively declines, reflecting a shift toward more efficient motor control. Similarly, Osu et al. (2002) found that trial-by-trial movement errors temporarily increased co-contraction in subsequent movements, whereas successful trials led to reduced co-contraction (7). This suggests that the CNS initially relies on impedance control to compensate for inaccuracies in the internal model and progressively reduces co-contraction as the model becomes more precise. Further supporting this, Heald et al. (2018) provided direct evidence that early muscle co-contraction facilitates motor adaptation (8). Participants pre-trained to co-contract (high impedance) during perturbations learned a novel force field faster than those who remained relaxed. Greater early co-contraction correlated with faster improvement, suggesting that early co-contraction is not just a side effect but an active strategy that accelerates internal model acquisition by reducing large movement errors. Beyond adaptation, Darainy & Ostry (2008) examined muscle coactivation after extensive training in stable dynamics and found that even after motor learning plateaued, co-contraction did not disappear (14). Instead, it remained a task-specific component of muscle activity, indicating that the CNS retains co-contraction as a control strategy for stability and precision, even in well-learned skills.

In contrast to prior studies relying on whole-muscle EMG, which could only infer co-contraction changes from bulk activation patterns (26), our single MU analysis provides direct evidence indicating that specific adjustments in MU recruitment and firing patterns drive alterations in co-contraction during adaptation. To generate the forces required for movement and counteract external perturbations, the CNS leverages two fundamental mechanisms at the MU level: MU recruitment (amplitude modulation) and rate coding (frequency modulation) (27). The recruitment process, which activates a larger number of MUs, especially those with higher activation thresholds, substantially increases the overall muscle force amplitude (16). This recruitment strategy is particularly critical when responding to novel perturbations that demand rapid and substantial increases in force or joint stiffness. Our findings demonstrate that during early adaptation, the CNS meets these demands by recruiting a larger pool of MUs across both agonist and antagonist muscles, effectively enhancing neuromuscular drive. In addition to MU recruitment, frequency modulation is essential for optimizing muscle force output and ensuring that the recruited MUs are precisely tuned to task demands (28). Consistent with Henneman’s size principle, the initial generation of low-to-moderate force levels within a muscle is predominantly achieved through MU recruitment, with subsequent fine-tuning of force achieved through firing rate modulation (16, 18). However, in conditions requiring rapid force adjustments and dynamic motor control, rate coding assumes an increasingly pivotal role. Our data indicate that during early adaptation, the CNS responds to unexpected perturbations by rapidly increasing the firing rates of recruited MUs, thereby generating quick corrective forces. As adaptation progresses and the perturbation becomes predictable, the CNS transitions to a feedforward control strategy, anticipating the force-field and activating the appropriate muscles in advance. Rather than relying on frequency modulation, the CNS refines MU recruitment, engaging additional triceps MUs, including larger, high-force units, which enables the required force to be achieved with stable, moderate firing rates.

In dynamic movements, increasing MU recruitment rather than increasing firing rates can be biomechanically advantageous. When more motor fibers share the load, each can operate in a favorable range, leading to smoother and more efficient force production. Conversely, increasing the firing rates of a few MUs to very high levels can result in force output nearing saturation. This may also lead to variability in force due to the limits of twitch fusion and fluctuations in firing rates. In our study, the CNS recruits additional MUs in the triceps during late adaptation to ensure the elbow extension force is distributed across multiple muscle fibers. This minimizes any potential tremor or oscillation that could arise if a single unit were operating at its maximal firing rate. Additionally, the continued increase in triceps MU amplitude suggests the recruitment of higher-threshold MUs to meet force demands in a viscous field as movement speed increases. This aligns with the principle that optimal MU recruitment ensures the necessary force output at a given movement velocity. From early to late adaptation, biceps MU activity remains stable, indicating that participants do not progressively increase antagonist involvement. As the force-field becomes familiar, biceps activity stabilizes, allowing the triceps to generate force unopposed. This reciprocal activation enhances mechanical efficiency, ensuring that all generated force contributes to movement or stabilization rather than being wasted in opposing muscle tension.

Our findings are consistent with established patterns observed in various contexts of motor skill learning. During the initial stages of coordination tasks, such as learning a new sport technique, novices often exhibit muscle overactivation, resulting in movements that are mechanically rigid and energetically inefficient. However, through practice and progressive skill acquisition, individuals learn to optimize muscle activation strategies, thereby achieving enhanced levels of task performance (29). Similarly, locomotor adaptation paradigms, such as split-belt treadmill walking, initially show elevated levels of muscle co-contraction, which decrease as gait becomes smoother and more efficient (30). These observations support the concept that the CNS’s strategy of refining MU recruitment is a fundamental principle in motor learning. Whether adapting to a controlled laboratory force field perturbation or mastering a new physical challenge in real-world scenarios, the brain initially prioritizes movement stability and successful task execution, before progressively streamlining MU activation patterns for improved neuromuscular efficiency.

While this study provides insights into MU changes during force-field perturbations, several limitations should be considered. First, the analysis focused exclusively on co-contraction between the biceps and triceps brachii, emphasizing elbow joint dynamics. Adaptation to new force environments likely requires coordinated activity across multiple joints, particularly the shoulder. Given that the task’s perturbation applied a velocity-dependent leftward force and a viscous force, shoulder muscles—such as the deltoid and rotator cuff—likely contributed to stabilizing and counteracting the force. Previous research has indicated that upper limb adaptation involves intermuscular coordination between shoulder and elbow muscles to generate precise corrective forces (31). Without direct measurements of shoulder muscle activity, these findings may not fully capture the broader neuromuscular adjustments occurring during adaptation. Future studies incorporating high-density EMG from the shoulder complex could provide a more comprehensive understanding of inter-joint coordination in motor adaptation. Second, the MU recordings represent a sample of active units rather than the entire motor pool. Although EMG signals were carefully decomposed to extract individual MU activity, it is possible that smaller or deeper units, which contribute to fine motor adjustments, were not captured. Additionally, tracking the same MUs throughout dynamic movements presents technical challenges, potentially introducing variability in assessing MU recruitment and firing rate changes. To address this, rigorous signal decomposition techniques were employed, and unit consistency across trials was verified, but some nuances of MU behavior might still have been missed. Despite these limitations, the findings provide important insights into the neuromuscular processes underlying dynamic force-field adaptation. The observed evolution of MU recruitment, amplitude, and firing rate suggests a transition from robust, error-driven control to refined, efficient force production. These results contribute to the understanding of motor adaptation processes at the MU level.

## Competing interests

The authors declare no competing interests

## Data availability statement

The data that support the findings of this study are available from the corresponding author upon reasonable request.

## Author Contributions

Y.R.E., S.B., and Y.L. designed research; Y.R.E. and S.B. performed research; Y.R.E., S.B., and Y.L. analyzed data; and Y.L., Y.R.E., and S.B. wrote the paper.

## Notes

### Competing Interest Statement

The authors have declared no competing interest.

